# Extracting nanoscale membrane morphology from single-molecule localizations

**DOI:** 10.1101/2023.01.26.525798

**Authors:** Zach Marin, Lukas A. Fuentes, Joerg Bewersdorf, David Baddeley

## Abstract

Membrane surface reconstruction at the nanometer scale is required for understanding mechanisms of subcellular shape change. This historically has been the domain of electron microscopy, but extraction of surfaces from specific labels is a difficult task in this imaging modality. Existing methods for extracting surfaces from fluorescence microscopy have poor resolution or require high-quality super-resolution data that is manually cleaned and curated. Here we present NanoWrap, a new method for extracting surfaces from generalized single-molecule localization microscopy (SMLM) data. This makes it possible to study the shape of specifically-labelled membraneous structures inside of cells. We validate NanoWrap using simulations and demonstrate its reconstruction capabilities on SMLM data of the endoplasmic reticulum and mitochondria. NanoWrap is implemented in the open-source Python Microscopy Environment.

**SIGNIFICANCE:** We introduce a novel tool for reconstruction of subcellular membrane surfaces from single-molecule localization microscopy data and use it to visualize and quantify local shape and membrane-membrane interactions. We benchmark its performance on simulated data and demonstrate its fidelity to experimental data.

## INTRODUCTION

Changes in cellular membrane shape are linked to viral replication, Alzheimer’s disease, heart disease and an abundance of other maladies (1–7). Understanding the morphological mechanisms behind these diseases requires imaging not only the locations of protein clusters causing structural changes in a membrane, but finding the underlying shape of the membrane at relevant size scales. Some membranous organelles, such as the endoplasmic reticulum (ER) and the Golgi, have diameters as small as ∼50 nm, requiring an image resolution of 25 nm or better to properly resolve structure (8, 9). Electron microscopy (EM) techniques have a resolution of 2 nm or better and are well-suited to imaging membranes. However, segmentation of membrane structures from EM images can be arduous (10, 11) and it can be difficult to label and identify specific proteins in EM samples. Immunolabeling with gold nanoparticles, the most common method for specifically labeling proteins in EM samples, requires fixation methods that often destroy cellular structures and suffers from relatively poor labelling efficiencies. It also appears in the same channel as all other cellular features, requiring non-trivial image post-processing to extract the locations of individual gold nanoparticles (12, 13).

Fluorescence microscopy techniques provide spectral separation of multiple fluorescent labels, allowing for easy identification of both membrane-associated and membrane-interacting proteins. However, conventional fluorescence microscopy techniques achieve a resolution no better than 250 nm and are therefore unable to visualize membrane curvature at true size scales. Single-molecule localization microscopy (SMLM) techniques, such as PALM, STORM, and PAINT, image the positions of proteins with ∼10 − 20 nm resolution (14). This is sufficient for imaging membrane structural changes of interest (15). In contrast to EM imaging techniques, which show a continuous membrane, SMLM yields a sparse and noisy point cloud of fluorophore locations, each with an uncertainty that depends on the brightness of the underlying blinking event. To visualize and quantify a membrane, it is necessary to interpolate a continuous surface through these positions.

In the fields of remote sensing (16) and 3D scanning (17), Screened Poisson Reconstruction (SPR) (18) is often used to extract surfaces from point clouds. SPR reconstruction is designed to follow point locations exactly, giving it high fidelity to collected positions. SPR has been applied to SMLM data (19), but this required extremely high quality data that was manually curated and pre-processed. Due to labelling inefficiencies and sampling, SMLM data often shows holes in large regions of a structure. To paper over these holes, SPR relies on the normals of surrounding point data to estimate the gradient of the original surface. Accurate normals are easily recovered in 3D scanning, but the high background in SMLM confounds standard estimation methods that rely on neighboring points to calculate normals. Fluorescent labels often bind to not only molecules of interest, but to other, non-specific targets in the sample. This and sample autofluorescence can generate spurious background localizations (20). The stochastic nature of SMLM imaging means each fluorescent molecule may blink multiple times thus appearing as multiple points, each in a slightly different spot. Adhering strictly to all of these points will not necessarily generate an accurate surface approximation of the underlying structure the point cloud represents.

In order to generate an accurate surface from general SMLM data, it is necessary to account for the localization precision of each point. By weighting each point’s influence on surface structure by its precision, a surface is allowed to move away from points with high uncertainty while still adhering to the data. This makes it possible to ignore contributions of poorly-localized spots arising from auto-fluorescence and non-specific binding. This is the key difference between our approach and other shrink-wrapping routines, which adhere strictly to or generate surfaces at a constant offset from input point clouds (21, 22) (see Figure S1). Shape priors have been used to smooth meshes sensibly over empty areas in point clouds, but these do not discriminate between background and structural points and standard loss functions that are robust to outliers, such as the one used in (23), do not account for the heavy background noise conditions seen in SMLM data. Zhao et al. created a surface fitting algorithm that takes pixel noise into account when generating surfaces from voxel-based images, and extended some of the paper’s maths to create an expression that uses localization uncertainty to fit SMLM data sets (24). However, the authors did not create an implementation for SMLM surface fitting. To our knowledge, no research group has demonstrated a general method for fitting surfaces to SMLM data that leverages information about localization uncertainties.

Here we present a novel algorithm, which we call NanoWrap, that creates constrained organelle surface representations from SMLM point clouds. NanoWrap incorporates localization uncertainty into its fitting routine and works for any 3D SMLM data set. Smoothing techniques are often applied to extracted surfaces to achieve reasonable shapes (25, 26), and such an approach is used in this algorithm. NanoWrap is implemented in PYthon Microscopy Environment Visualize^1^ for ease-of-use, integration with additional SMLM acquisition and analysis techniques, and the ability to pass localizations acquired via a user’s microscope and software of choice to the algorithm (27).

## MATERIALS AND METHODS

### Initial / starting mesh

The major steps of NanoWrap are sketched in Figure 1. The algorithm takes an initial, coarse surface that loosely approximates a point cloud and iteratively refines the structure under point fidelity and curvature constraints.

**Figure 1.**
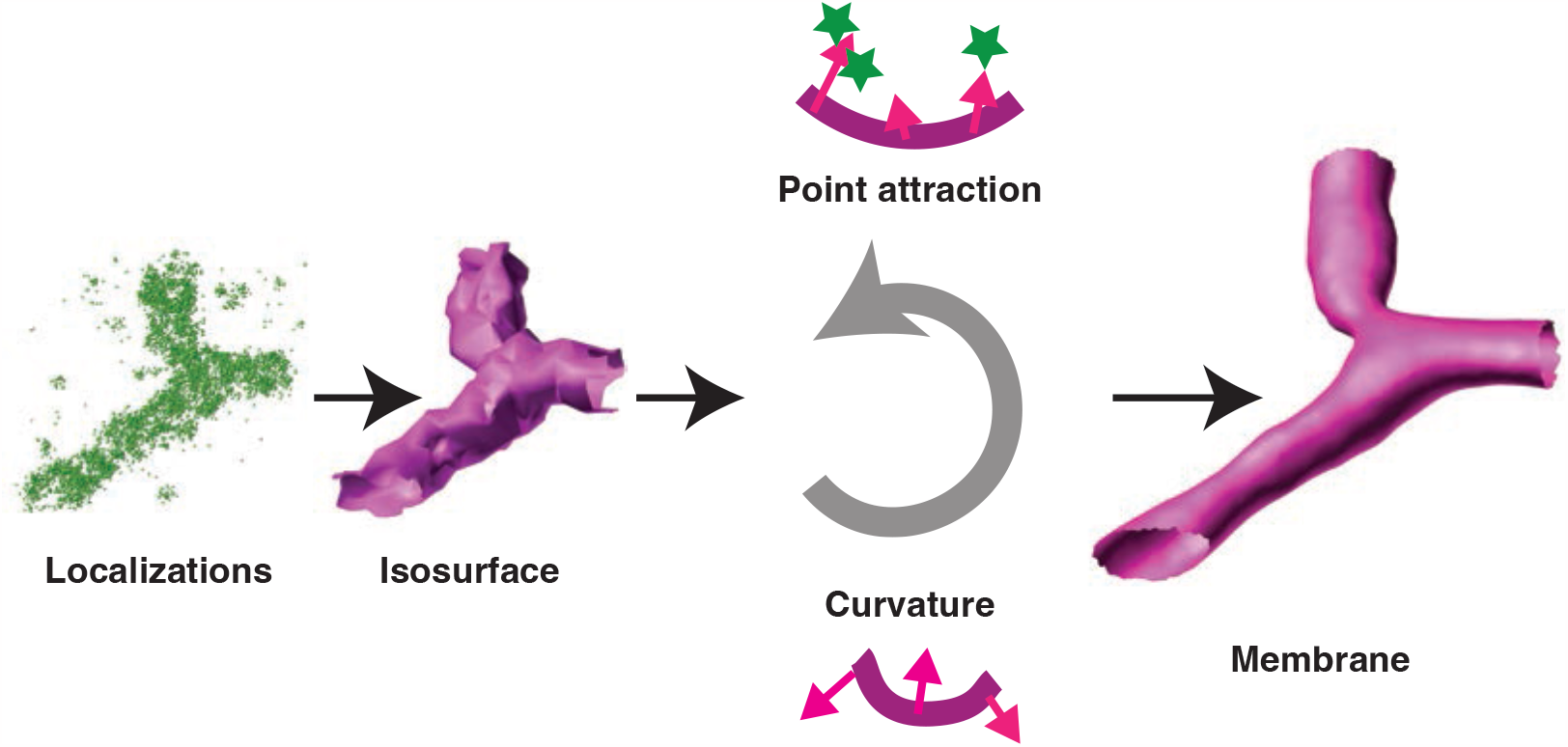
: A flow diagram representing the major steps of NanoWrap. Localization data is first approximated by a coarse, density-based isosurface. This surface is moved toward the localizations subject to a curvature force constraint. The pipeline runs iteratively until stopping criteria are met. The result is a membrane approximation of the underlying continuous structure sampled by the input localizations.

First, a set of single-molecule localizations are placed in a sparse octree data structure (28). The octree is truncated at a given minimum number of localizations per octree cell (equivalent to a minimum signal-to-noise ratio—see (29)). This has the effect of dividing the volume into cubic cells with sizes that adapt to local point density. Cells will be large in areas with few localizations, and small in areas which are localization dense. Cells containing fewer than the minimum number of points are not stored. The result is a volumetric data structure that contains the same information as a regularly sampled grid, but requires significantly less memory. The density of localizations in each cell is calculated. The Dual Marching Cubes algorithm is run on these cells with a given threshold density (30). The result is a manifold triangular mesh that separates high from low density areas (27). Because of SMLM labelling is sparse, a density-based isosurface can only recover a crude representation of the underlying structure. To avoid issues with over-segmentation, the threshold for this starting estimate should be chosen such that the resulting surface lies outside the true membrane surface (i.e. using a lower threshold value than would be used if trying to estimate the surface directly with an isosurface). The following steps will then pull the surface in onto the true membrane location.

### Mesh optimization

The resulting mesh is refined to achieve high fidelity to the localization input data under a curvature constraint. This is expressed mathematically as a minimization problem:

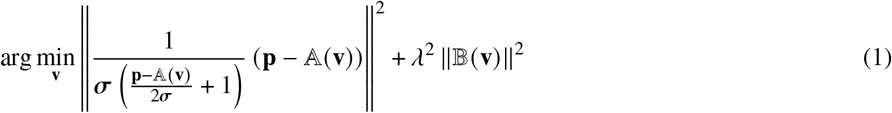

where **p** is a vector containing localization positions, **v** is a vector containing mesh vertex positions, the **p** -𝔸 (**v**) term represents the distances between each localization and the mesh, 𝔹 (**v**) encodes a curvature penalty, λ is a constant controlling the relative weighting of point fidelity and curvature terms and *σ* is a vector containing localization uncertainties. In microscopic imaging, the localisation error ellipse *σ* can generally be assumed to be aligned with the Cartesian axes (using the common convention that x and y are the camera axes, and z is along the optical axis of the microscope) and it is possible to consider errors in x, y, and z independently. If our method were applied to methodologies (e.g. LIDAR) where this assumption cannot be made, the method would need to be adapted to use a covariance matrix.

The above equation is iteratively solved using a conjugate gradient method (31). If, after refinement, the mesh does not change shape significantly as compared to its shape in the previous iteration, or if the given maximum number of iterations is reached, the algorithm terminates and the resulting surface is presented as an approximation of the underlying structure. Otherwise, the surface is passed to the mesh refinement step and all subsequent steps repeat.

### Point fidelity term

#### Notation

**p** the *N* × 3 array of fluorophore localizations

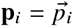 the *i*th localization (1 × 3)

σ *N*×3 array of the uncertainty of the localizations

σ_*i*_ the uncertainty of the 8th localization (1 × 3)

**V** the *M*× 3 array of mesh vertices

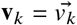 the *k*th mesh vertex (1 × 3)

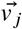 the *j*th vertex of a given mesh face

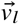 the *l*th vertex neighbour of a given vertex

The point attraction term seeks to minimize the distance between each localization and the surface. For each localization, we approximate its distance to the surface as the distance to a proxy point formed by a linear combination of the three vertex positions that define its closest surface face. The weightings used in the linear combination are computed as follows, based on the inverse distance from a localization 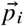 to each vertex 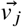 in its nearest face.

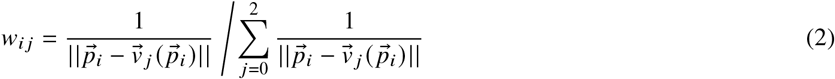

The position of the proxy point is then calculated as

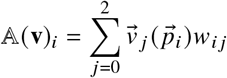

giving the distance metric:

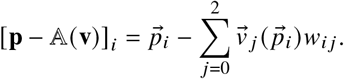

This metric is asymptotically equal to the true distance to the surface at small distances, but trends to the distance between the localization and the center of its nearest face (all vertices weighted equally) at large distances. This behaviour was deliberately chosen to ensure sensible updating of vertex positions—for a localization close to the surface we want it to mostly pull on the closest vertex, whereas a point that is far away (compared to the face edge length) should pull equally on all vertices of the face.

This distance metric is weighted by

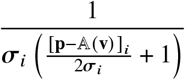

as shown in Equation 1. The 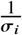 term ensures distances near a localization produce minimal cost. The remaining fraction de-weights localizations that are particularly far away, while still letting them exert influence on the surface. This ensures the surface can shift to accurately fit all points in a point cloud if necessary, but places priority on moving toward the centroid of its nearby points first. Since localizations are Gaussian-distributed, they should exercise most of their influence within 2*σ*_*i*_ of their position.

### Curvature term

When minimizing curvature at a vertex, there are multiple possible formulas to choose from. One of the simplest is the Laplacian discretization of the Canham-Helfrich bending energy (32),

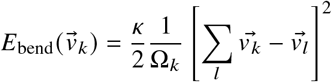

where 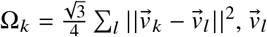 is the *l*th neighbor of vertex 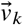, and *κ* is the stiffness coefficient for the lipid composition of the membrane. The local bending energy at vertex 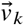 can then found by minimizing

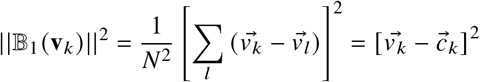

or the squared distance between a vertex and the centroid of its neighbours 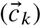 . This formulation, however, leads to a shrinking of the membrane, with a surface constrained entirely by curvature eventually collapsing to an infinitely small sphere (a valid, but trivial solution to the bending energy minimization problem). As a result, balancing this curvature term with a point-attraction term will always lead to a surface approximation that lies inside the true surface. In practice, the shrinking effect is sufficiently strong that a curvature weighting (λ) low enough for the point attraction to prevent excess shrinkage will lead to an excessively rough surface.

An alternative, area-preserving approach to curvature minimization (𝔹_2_) is to penalise the distance between a vertex and a location on the surface of a sphere fit to the vertex’s neighboring vertices. 𝔹_2_ can be approximated as

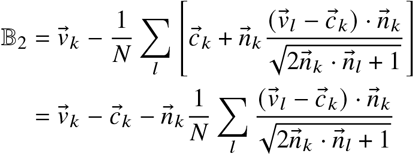

where 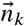 and 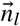 are the normals of vertex *k* and its *l*th neighbour. This parameterization works when the surface is well-constrained by localization data, leading to a smooth surface with good affinity to the localizations, but can result in large, static “blebs” in areas where the starting estimate was poor and localizations are especially sparse (see Figure S2). Our empirical solution is to use 𝔹_2_ where the influence of localization data is high, smoothly transitioning to the area minimizing approach, 𝔹_1_, as influence decreases. This gives 𝔹 the following form:

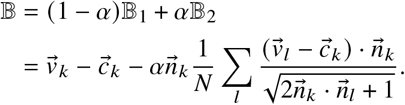

The empirical value for alpha is

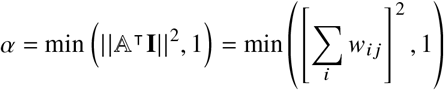

where 𝔸 is the operator that generates point-fidelity proxy points and *w*_*ij*_ is as defined in equation 2 above.

### Mesh refinement

The starting mesh we derive from density thresholding is relatively coarse (consisting of triangular cells which are large compared to organelle curvature). The initial mesh will typically also erroneously connect parts of the structure that should be separate (undersegmentation) -a desirable attribute as it is easier to separate areas which were erroneously joined than to join areas which were erroneously separated. As the optimization progresses, it will also expand or contract areas of the mesh unevenly, leading to a wide variation in mesh cell size. To resolve these issues, we interleave optimization steps (described above) with operations which manipulate the mesh, performing “remeshing” operations which lead to a more finely sampled mesh and ensure that the mesh is well formed for subsequent numerical operations (33), and “neck removal” operations, which cut the necks which form as the mesh wraps around areas that were erroneously joined in the starting mesh. These operations are performed every 5 iterations. Results are not strongly dependent on this frequency (see Figure S3), although it should be regular enough to keep the mesh edge lengths fairly constant as parts of the mesh pull in. It is preferable to choose the total iteration number such that a few shrink-wrapping iterations are performed after the final remesh.

Remeshing (33) consists of a number of operations—splitting long edges, collapsing short edges, “flipping” edges where too many are incident on a vertex, and regularly spacing vertices along the surface. Together, these remeshing operations result in a well-formed mesh where the edge lengths are roughly constant, the number of edges incident on a single vertex is roughly constant, and the triangles are roughly equilateral. By adjusting the target edge length in the remeshing operation we can adjust the sampling of the mesh. Starting from the mean edge length of the starting mesh, we decrease the edge length linearly toward 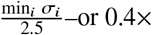 the minimum localization precision; a value that lets us adequately sample the smallest resolvable features in our localization dataset. Alternatively, a minimum edge length can be set by the user. This edge length should be sufficiently small to sample the curvature expected in the structure. In practice, a lower limit of 5 nm works well for the structures shown in this paper. The linear reduction of edge length during the course of optimisation ensures the iterative fitting makes large adjustments early in the fitting and fits detailed features later on.

To remove necks, vertices with unphysically high negative Gaussian curvature are deleted, and resulting holes in the mesh are stitched with fresh triangles. This operation removes false, thin necks in between portions of the surface. This operation is performed in regular intervals throughout our iterative fitting process.

### Simulation and quality evaluation

To evaluate the fidelity and accuracy of our method, and to compare with the results obtained using SPR we applied both methods to simulated data for which the ground truth surface was known. SMLM point clouds were simulated from a theoretical figure eight, defined by a signed-distance function (see Supplementary Information). Simulations varied point-cloud density and number of background localizations. We did not explicitly vary localization precision as, being mathematically equivalent to simple scaling of the structure, this should effect both methods equally.

Surfaces were fit to simulated point clouds and then compared to the theoretical structure giving rise to these point clouds. When making this comparison we must consider two types of error 1) the distance from the true surface to the reconstructed surface and 2) the distance from the reconstructed surface to the true surface. Although these two metrics might seem redundant at first glance, they are not. It is possible for a surface to appear good under metric 1 whilst having bad performance under metric 2. An example of this is a structure that mostly closely follows the true surface, but also has “blebs” or extrusions away from the true surface. Because the distance in metric 1 just considers the parts of the reconstruction which are closest to the true structure, “blebs” and extrusions are not penalised and metric 1 returns a small distance. Similarly, a reconstruction which follows part of the ground truth correctly, but is truncated such that it does not extend into all areas of the ground truth, can score well on metric 2. A good reconstruction minimizes both of these metrics.

The distance between surfaces was calculated numerically as follows; a set of of noise-free verification points were simulated exactly on the surface of the theoretical structure, and a set of noise-free points were simulated on the fit surface. The mean squared distance from the verification point set to its nearest neighbors in the mesh point set was computed as quality metric *Q*_1_. The mean squared distance from the mesh point set to its nearest neighbors in the verification point set was computed as quality metric *Q*_2_. Mesh quality was scored as a combination of the two error types:

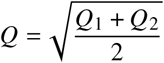

where *Q* is the root-mean-square error, representing the average distance from the mesh to the theoretical structure.

### Cell culture and sample preparation

U-2 OS cells (HTB-96; ATCC; Lot: 70008732) were grown in McCoy’s 5A medium (16600-082; Gibco) supplemented with 10% FBS (10438-026; Gibco). These cells were subcultured with 0.05% Trypsin (Gibco).

Immunofluorescence samples were generally prepared as follows. Approximately 1 million cells in 90 µL were transfected via electroporation with about 10 µg of plasmid with a Super Electroporator NEPA21 Type II (Nepa Gene). The cells were then seeded onto coverslips that were, unless otherwise noted, treated in an ozone chamber for 30 minutes. Cells were chemically fixed the following day with 3% paraformaldehyde (15710; Electron Microscopy Sciences) and 0.1% glutaraldehyde (16019; Electron Microscopy Sciences) for 15 minutes at room temperature while gently rocking. The glutaraldehyde fixation was quenched by washing the samples with 0.1% sodium borohydride in 1 PBS for 7 minutes followed by washing with 100 mM glycine in 1 PBS for 10 minutes. Samples were then rinsed three times with 1 PBS followed by a 3-minute incubation at room temperature with permeabilization buffer (0.3% IGEPAL CA-630 + 0.05% Triton X-100 + 0.1% (w/v) BSA in 1 PBS) and another three rinses with 1 PBS. Samples were blocked for 1 hour at room temperature in block buffer (0.05% IGEPAL CA-630 + 0.05% Triton X-100 + 5% normal goat serum in 1 PBS), followed by an overnight incubation with the primary antibody diluted in block buffer at 4o C while gently rocking. The following day, the samples were washed three times, for 5 minutes each wash, with wash buffer (0.05% IGEPAL CA-630 + 0.05% Triton X-100 + 0.2% (w/v) BSA in 1 PBS), incubated with secondary antibodies diluted in block buffer for 1 hour at room temperature while gently rocking, washed three more times, for 5 minutes each wash, with wash buffer, and finally rinsed three times with 1 PBS. For astigmatic/4Pi-DNA-PAINT imaging of overexpressed mCherry-Sec61*β* and 4Pi-DNA-PAINT imaging of TOMM20-mCherry, samples were immunolabeled with rabbit anti-mCherry primary (ab167453; Abcam) diluted 1:500 in block buffer and rabbit anti-TOMM20 primary (sc-11415; Santa Cruz Biotechnology) diluted 1:500 in block buffer, respectively. Both samples were then labeled with an oligonucleotide-conjugated goat anti-rabbit IgG secondary antibody (115-005-146; Jackson ImmunoResearch) diluted 1:200 in block buffer, as described previously (34). For two-color 4Pi-STORM imaging of overexpressed mCherry-Sec61*β* and endogenously expressed TOMM20, samples were labeled with a combo of two mouse anti-mCherry primaries (GTX630195 and GTX630189; GeneTex) each diluted 1:250 in block buffer and rabbit anti-TOMM20 primary (see above) diluted 1:500 in block buffer. They were then labeled with a goat anti-rabbit secondary conjugated to AF647 (20812; Biotium) diluted 1:1000 in block buffer and a goat anti-mouse secondary (A21245; Invitrogen) with a single conjugated CF660C dye on each secondary antibody diluted 1:1000 in block buffer. The mCherry-Sec61*β* plasmid was acquired from Addgene (49155).

### Microscopy

4Pi-SMS two-color samples of TOMM20 and mCherry-Sec61*β* were prepared and imaged using ratiometric dSTORM as described previously (35). 4Pi-SMS one-color DNA-PAINT samples were imaged on the same custom microscope, but with fluorogenic DNA-PAINT (see below). Astigmatic data was collected using a custom-built microscope described previously (36) with the only filter used being a bandpass filter (FF01-694/SP; Semrock). Astigmatism was implemented by adding a cylindrical lens to the fluorescence light path.

All DNA-PAINT data was collected using the fluorogenic DNA-PAINT method described previously (34). Briefly, samples were imaged using the following imager probe containing a dye and a quencher: 5’ -Cy3B -AAGAAGTAAAGGGAG -BHQ2 -3’. This imager probe was diluted to 10 nM for both astigmatic and 4Pi imaging of mCherry-Sec61*β* and 1 nM for 4Pi-SMS imaging of TOMM20-mCherry in a high ionic-strength PBS-based buffer (1 PBS, 500 mM NaCl, 20 mM Na_2_SO_3_, and 1 mM Trolox, pH 7.3-7.5).

## RESULTS AND DISCUSSION

### Validation on simulated SMLM datasets

A three-dimensional figure eight (two toruses touching) was simulated across a range of localization density and background values chosen to reflect the ranges we see in typical experiments (see Supplementary Information). As both SPR and NanoWrap have a number of user settable hyper-parameters, a fair comparison of the methods requires a systematic search of the parameter space to avoid the possibility that we simply choose a disadvantageous SPR parameterization. For each condition, we therefore performed a grid-search of the feasible parameter space for both SPR and NanoWrap. The resulting meshes were scored as described in Quality evaluation. For each condition, the mesh with the lowest RMS error (*Q*, see Quality evaluation) from each method was selected for comparison. This eliminates the possibility of user bias in parameter selection. The results are shown in Figure 2.

**Figure 2.**
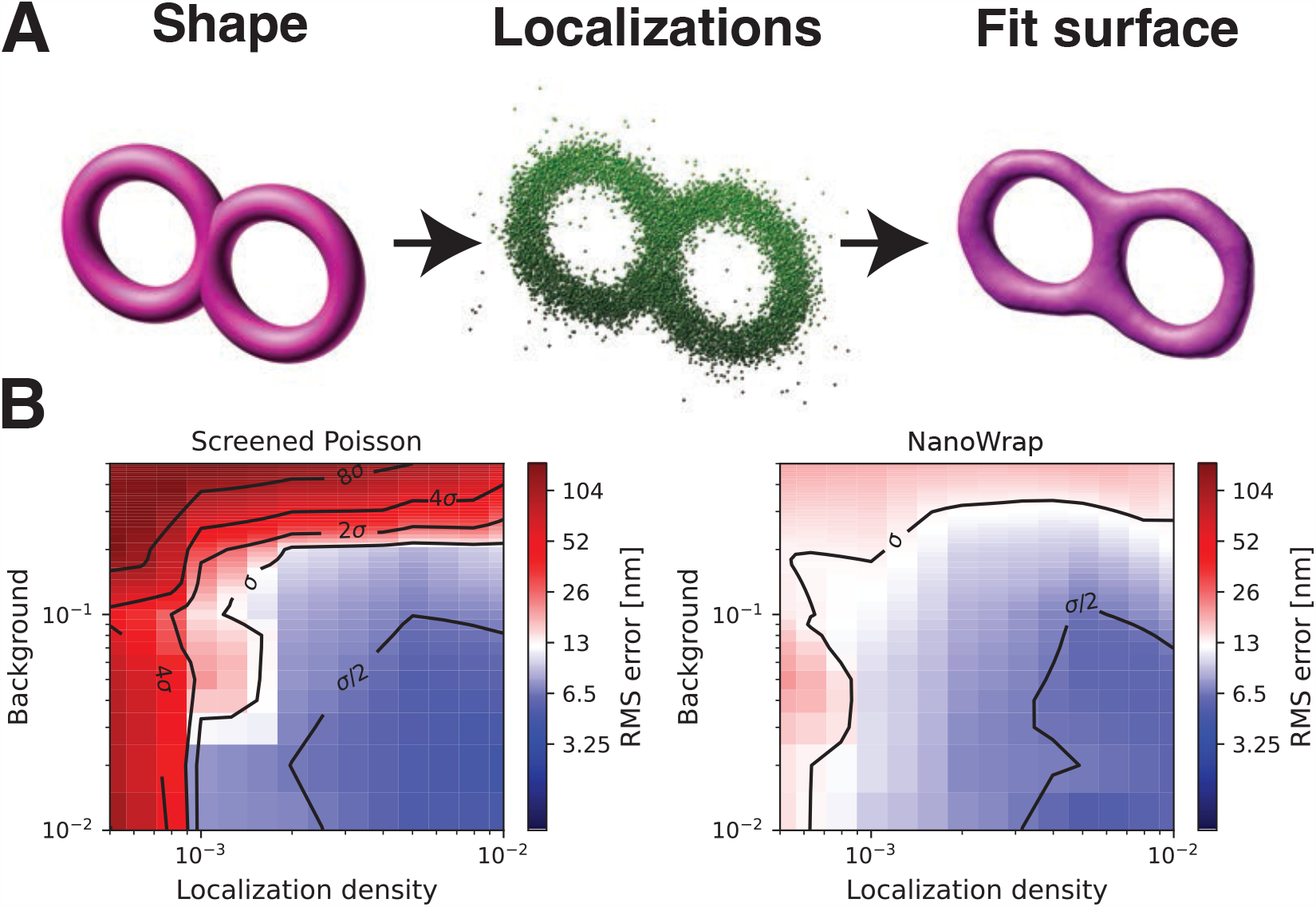
: A comparison of SPR and NanoWrap on simulated point clouds of a 3D figure 8, which is approximately 800 nm in diameter and 100 nm thick. **A**, Our simulation method of going from a signed distance function to a set of localizations to a fit surface. **B**, Heat map plots of Screened Poisson Reconstruction and NanoWrap as functions of simulated localization density and background. For reference, the experimental SMLM data shown in figures 3 and 4 had a median localization density of approximately 6 × 10^−2^nm^-2^ and background ration of 2 ×10^−1^.

Both NanoWrap and SPR work well when there is a high density of points and low background, and are capable of delivering a RMS error between ground truth and fitted surfaces which is less than the localization precision (*σ*). The combination of low background and high density is rare, however, in experimental data. As density decreases and background increases, SPR has a hard time fitting an accurate surface. In contrast, NanoWrap yields surface errors which are better than the localization precision (*σ*) and can accurately estimate curvatures over a wide range of densities and background. Performance also degrades only slowly outside this space, with acceptable performance even at the very low localization densities that might be seen in live-cell SMLM.

To investigate the accuracy of our curvature estimates, we simulated a tapered cylinder varying in radius from 30 nm to 200 nm over a 2000 nm length (Figure S4 and Figure S5). As might be expected, the accuracy of curvature estimation depends on both localisation density and precision, becoming unreliable at very low densities and poor localisation precision. Based on these simulations, we are likely to under-estimate curvature when the true curvature radius is less than approximately 3 times the localisation precision or about twice the median distance between localizations.

The sensitivity of our algorithm to parameter choice is discussed in detail in the Supplementary Material (Supplementary Note S.3 and Supplementary Figure S6). Briefly, curvature weight is the most important user-settable parameter and is used to balance the point fidelity and curvature terms. The optimal value of this parameter increases as point density and background increase. The corresponding minimum in surface error, however, is broad and small changes in the parameter value do not have large effects on surface quality.

### Application to experimental SMLM data

We initially tested our NanoWrap algorithm on data acquired using 4Pi-SMS microscopy (35, 37) as this provides high precision isotropic localization sufficient to capture the finest details of small intracellular organelles such as the endoplasmic reticulum (ER). When applied to this data, NanoWrap results in an approximation of the ER surface (Figure 3) which is consistent with both the observed Sec61*β* localizations and our knowledge of ER structure. Cross-sections of the data allow us to visually confirm that the surface approximation is well matched to the localization point-cloud data (Figure 3E-G). Quantification of ER tubule diameter from the surface reconstruction is in good agreement with previously reported values (38) (see Figure S9).

**Figure 3.**
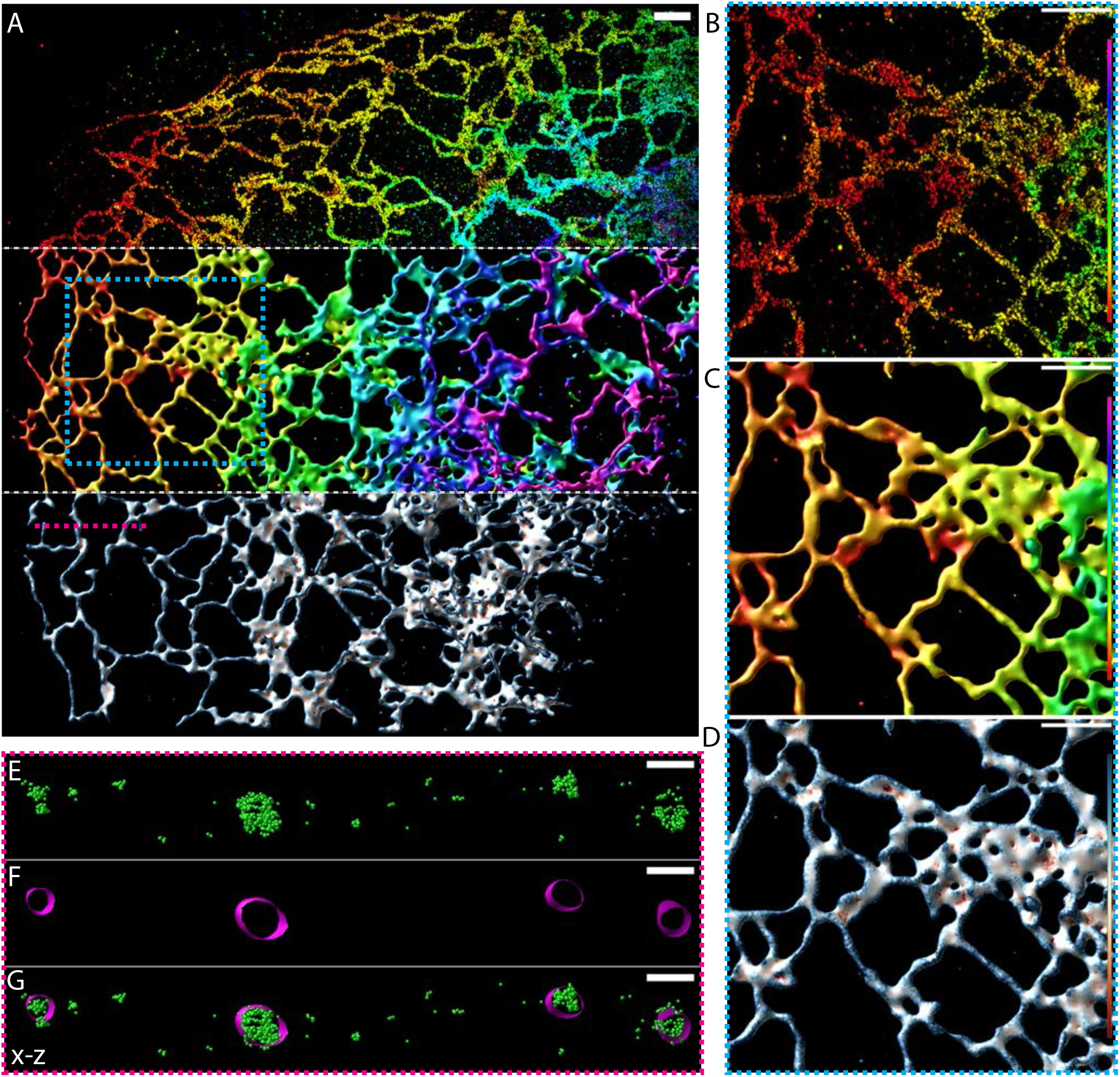
: **A**, 4Pi fluorogenic DNA-PAINT data of overexpressed mCherry-Sec61*β*,an endoplasmic reticulum membrane protein ; Top: Point-cloud data displayed as 10 nm point sprites with alpha set to 0.5 and colored by each point’s position in z according to the lookup table described in B/C; Middle: NanoWrap surface created based on the point-cloud data colored by each vertex’s position in z according to the lookup table described in B/C; Bottom: The same surface, but colored by the mean curvature at each vertex according to the lookup table described in D. **B-D**, ROI shown by the hashed cyan box in A displayed in the three different ways described in A. B-C lookup table (from bottom to top): 0 to 800 nm. D lookup table: -0.01 to 0.01 nm^-1^. **E-G**, Cross section in x-z shown by the hashed magenta line in A. **E**, Point cloud displayed as 10 nm green spheres. **F**, Surface displayed in magenta. **G**, Point cloud and surface displayed together. Scale bars are 1 µm **(A-D)** and 200 nm **(E-G)**.

Although advanced localization techniques such as the combination of 4Pi-SMS and DNA-PAINT (or, alternatively, MIN-FLUX (39)) are likely to yield the highest quality reconstructions, these techniques are not as widely available as more accessible methods such as astigmatic 3D DNA-PAINT. To demonstrate that the algorithm is not restricted to the ER and can be used across a broader range of instrumentation and labelling approaches, surfaces were additionally generated for mitochondria and using 4Pi two-color dSTORM as well as astigmatic 3D DNA-PAINT, with the results shown in Figure 4. Notably, a good reconstruction was still possible from astigmatic data despite the poorer axial localization precision when compared with 4Pi-SMS (Figure 4 G-I). This can be attributed to the fact that our surface fitting approach takes this anisotropic error into account when weighting the attraction force and finds the average surface that passes through multiple nearby localizations (even if the scatter of points in z is larger, poorer axial precision will not change their mean position).

**Figure 4.**
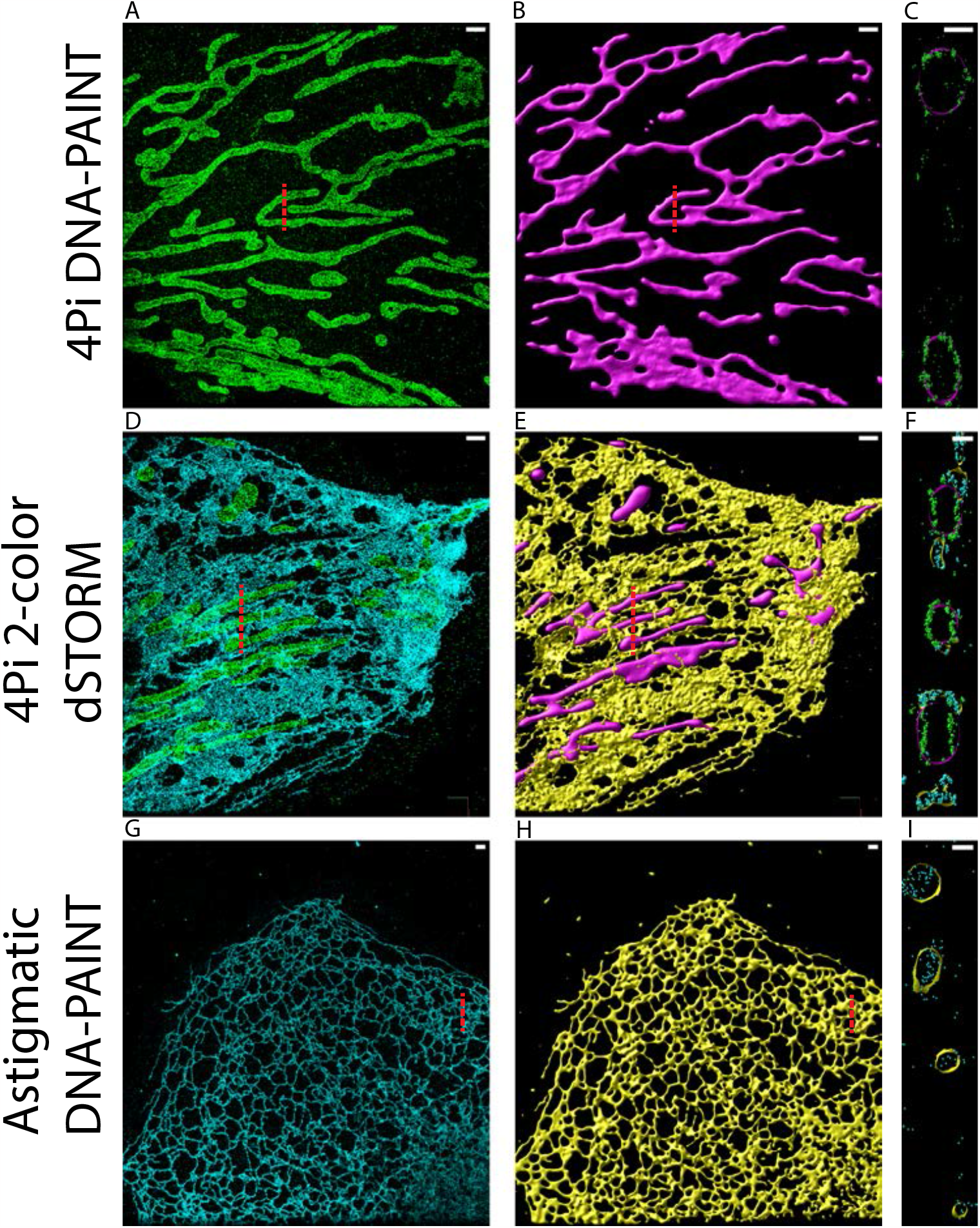
**Application of NanoWrap on data from varying SMLM imaging modes. A-C**, 4Pi fluorogenic DNA-PAINT localizations from outer mitochondrial membrane protein TOMM20 (green) and their resulting surface (magenta). **D-F**, Two-color 4Pi dSTORM localizations of TOMM20 (green) and overexpressed endoplasmic reticulum membrane protein Sec61*β* (cyan) and their resulting surfaces (magenta and yellow, respectively). **G-I** Astigmatic 3D fluorogenic DNA-PAINT localizations from overexpressed Sec61*β* (cyan) and their resulting surface (yellow). **A/D/G**, x-y view of localized point cloud. **B/E/H**, x-y view of resulting surface. **C/F/I**, x-z view of surfaces and points overlaid. Scale bars are 1 µm **(A**,**B**,**D**,**E**,**G**,**H)** and 200 nm **(C**,**F**,**I)**.

To achieve accurate curvature estimates using NanoWrap, localization precision must be smaller than the radius of curvature of the feature of interest (Figure S5). Given sufficient density and localization precision, NanoWrap is able to follow this curvature profile of a tapered cylinder smoothly (Figure S4). At too low a density, the curvature is slightly under-estimated. To reconstruct the ER (which has typical tubule diameters of around 100 nm) (38) we therefore recommend a median axial localization precision better than 30 nm (easily achievable using astigmatic 3D DNA-PAINT, but challenging for non-4Pi dSTORM), but predict that acceptable results will be able to be obtained on larger structures (e.g. mitochondria with a diameter of 0.5-1 µm) for all standard 3D localisation microscopy modalities. As implemented, NanoWrap will generally recover holes bigger than around three times the localisation precision, but will occasionally miss small holes or fenestrations in sheet like structures that might be discernible when examining the localisation cloud by eye (see also Supplementary note S.3 and Supplementary figure S8). This is both because the reliable detection of small holes is a surprisingly tricky computational problem, and because of a deliberate design choice to bias the starting surface this way so as to have a predictable failure mode and to limit the types of topological errors that must be handled in subsequent steps. Attempting to be bias free leads to fragmentation of structures and the generation of spurious holes before all true holes are reliably detected. Although not worse than other available algorithms, improving the topological accuracy of NanoWrap is a promising target for future research.

The surface approximations we create allow exciting new opportunities for quantitative analysis, permitting, for example, the measurement of curvature at each point on the surface (Figure 3A and D). In two-color data, each color channel can be fit independently (Figure 4 D-F) to yield relationships such as distance between surfaces to be calculated and visualized on the surfaces themselves (see Figure S10). This particular use has the potential to provide information about the location of membrane contact sites between the ER and other organelles that exhibit exceptionally small distances, 10 to 30 nm, between membranes (40). Analogous approaches can be used to measure distances to individual localizations in a second channel allowing the study of proteins which are partially or fully cytosolic.

## CONCLUSION

Examining the interplay of membrane surfaces and proteins is critical to understanding cellular function (41). The surface fitting algorithm described in this paper provides a new way for researchers to segment and quantify membrane structure from single-molecule localization microscopy data, and to subsequently study membrane-membrane and membrane-protein interactions at biophysically-relevant size scales. The algorithm and its resulting structures are biophysically-informed: localizations are fit based on expected fluorophore blinking behavior and labelling inefficiencies, and the curvature smoothing is inspired by the Canham-Helfrich bending energy functional, an established model for biological membranes (32, 42). This incorporation of prior information to achieve better reconstructions is a natural next step in biological membrane surface approximation and, more generally, to the analysis of noisy imaging data as a whole. It complements exciting new machine learning (ML) techniques for segmentation and analysis (11, 43, 44). In contrast with ML approaches, it does not require large volumes of manually annotated training data, and has a clear basis in biophysical theory.

The mathematics behind NanoWrap are not intrinsically limited to SMLM, and the ability to deal with noisy data, incomplete labelling, and anisotropic resolution should be a significant benefit for other high-resolution modalities (e.g. stimulated emission depletion (STED) and electron-microscopy). We look forward to adapting the method to these domains. A particularly exciting potential application of our technique is to live-cell super-resolution, which is heavily constrained by the achievable localisation density (SMLM) (45) or photon-count (STED) (46). The ability of NanoWrap to recover accurate organelle surfaces from very low localization densities (Figure 2) suggests an approximately 10 to 20-fold reduction in the required localization density. When combined with multi-emitter approaches, this should be enough to make high-quality live super-resolution imaging of dynamic organelles possible.

In summary, NanoWrap outperforms previously demonstrated methods of SMLM membrane estimation and enables high fidelity membrane reconstructions across a wide range of localization densities and backgrounds. For ease-of-use, shareability and adaptability, it is packaged as open-source software and is accessible via an interactive GUI. It is sufficiently fast and memory-efficient to be used on standard lab computers (e.g. mid-range laptops). NanoWrap is available for download as a PYME plugin at https://github.com/python-microscopy/ch-shrinkwrap.

## AUTHOR CONTRIBUTIONS

ZM and DB designed and implemented the algorithm. LF tested the algorithm, prepared samples, imaged the biological data and created the surface reconstructions of the biological data. JB provided critical feedback on the algorithm and the biological test case. All authors wrote the manuscript.

## ACKNOWLEDGMENTS

The authors wish to thank Michael Murrell, Megan King, Andrew Barentine, Yongdeng Zhang, Lena Schroeder, Florian Schueder and Frederic Pincet for helpful discussions.

We acknowledge funding from the Wellcome Trust (203285/B/16/Z) and NIH (R01GM118486, T32GM007223 and T32EB019941). This work is solely the responsibility of the authors and does not necessarily represent the official views of the NIH.

## Footnotes

1 https://python-microscopy.org

