## Supplementary Material for "Extracting nanoscale membrane morphology from single-molecule localizations"

### S.1 Point cloud simulation

In order to demonstrate the effectiveness of the algorithm, it was necessary to generate biologically-motivated test structures. To do this, we established a constructive solid geometry library where each primitive is represented as a signed distance function (SDF). An SDF describes an object by taking a point location as input and returning the signed distance between the point and the surface of the object. An example SDF is that of a sphere of radius $R \in \mathbb{R}$, $d(p) = ||p|| - R$, where $p \in \mathbb{R}^3$. $d(p) = 0$ indicates that point $p$ is on the surface of the sphere. Points located within the sphere have a negative distance to the sphere surface and points outside of the sphere have a positive distance to the sphere surface.

Points were generated from a test structure using Monte-Carlo sampling of an octree of the structure's SDF. The octree subdivides space that straddles where the SDF goes to zero until a fixed sampling rate $dx$ is reached. The center of each octree box with an absolute SDF value of less than $dx$ is kept with a probability of $p$ and rejected otherwise. This results in a point set of size $S$.

Next, non-specific localizations are simulated. For a given noise fraction $n$, $J = \frac{nS}{1-n}$ additional points are generated with uniform randomness over the bounding box of the point set. This ensures $J/T = n$ for $T = J + S$ total points.

To simulate localization precision, each point $i$ in the resulting point set is jittered by sampling values from a Gaussian with mean $\mu_i$ equivalent to the location of point $i \in 0..T-1$ and standard deviation $\sigma_{i,e}$ for $e \in x, y, z$. Up to $N$ samples are generated per point. The total point set is then clipped to size $T$ via random uniform selection of points.

The uncertainty $\sigma_{i,e}$ is calculated by simulating an exponential distribution with a mean value of $\rho$, corresponding to the average photon count of a simulated localization. This distribution is clipped to only contain values greater than a background photon count $b$, simulating the noise floor for data collection. For the remaining counts, the first $S$ are selected and

$$\sigma_{i,e} = \frac{r_e}{2.355\sqrt{\rho_i}}$$

where $r_e$ is the resolution of the simulated system along axis $e$.

### S.2 NanoWrap parameters

To generate the parameter space for evaluation of SPR meshes derived from simulated data in Figure 2, we followed recommendations in (18). For SPR we estimated normals from 10, 30, 50 and 100 nearest neighbors, set the smoothing parameter $\alpha = 0, 1, 2$ and 4, and set the samples-per-node to 1.5, 5, 10, 20 and 30. We used an octree depth of 8, 8 Gauss-Seidel relaxations, and a scale factor of 1.1.

For our method, we compared maximum number of iterations 19 and 39, a variety of starting threshold densities, $5 \times 10^{-6}$, $1 \times 10^{-5}$, $2 \times 10^{-5}$, $5 \times 10^{-5}$, $1 \times 10^{-4}$, $2 \times 10^{-4}$, $4 \times 10^{-4}$, $1 \times 10^{-3}$, $2 \times 10^{-3}$, and $\lambda = 10, 15, 25$. We remeshed every 5 iterations.

| | Max iters | Curvature weight | Remesh frequency | Neck first iter | Punch frequency | Kc | Minimum edge length | Smooth curvature | Truncate at |
|---|---|---|---|---|---|---|---|---|---|
| Figure 3 | 19 | 15 | 5 | 0 | 0 | 1.0 | 5.0 | True | 1000 |
| Figure 4 A-C | 39 | 10 | 5 | 0 | 0 | 1.0 | 5.0 | True | 1000 |
| Figure 4 D-F (ER) | 19 | 10 | 5 | 0 | 0 | 1.0 | 5.0 | True | 1000 |
| Figure 4 D-F (Mito) | 69 | 30 | 5 | 0 | 0 | 1.0 | 5.0 | True | 1000 |
| Figure 4 G-I | 15 | 10 | 5 | 0 | 0 | 1.0 | 5.0 | True | 1000 |

Table S1: Parameters used for iterative surface fitting of experimental data in this paper.

### S.3 Parameter sensitivity

The search of the parameter space used to quantify RMS surface error (Figure 2) also permits an analysis of the sensitivity of that error to parameter choices. As the starting surface is an isosurface on density, the optimal choice of the threshold parameter for this is expected to correlate to labeling density. In practice, however, as long as the starting estimate is not too far off, the subsequent optimisation steps will ensure good surface quality. Because the number of localizations pulling on a mesh face also correlates with density, the curvature weight required to balance the point attraction force is also expected to increase with increasing density. Figure S6 shows this is the case, although the minimum in RMS error is fairly broad indicating that the algorithm is not overly sensitive to the exact curvature weight. Increasing the background at a constant density shifts the optimal curvature weight to slightly higher values, but is not as significant as a change in density. The underlying curvature of the structure itself has only a very slight effect on the optimal curvature weight (Figure S7) the localization errors are already considered in the algorithm we do not expect a strong change in the optimal curvature weight with changing localization precision.

Unlike the RMS surface error, which can tolerate quite large variations in the starting surface, estimating the correct genus (number of holes) depends strongly on the threshold used to generate the starting surface. This is consistent with other shrink-wrapping approaches, which rely on manual thresholds to set hole size (21, 22). We attempt to address this automatically in NanoWrap, but to avoid obvious misrepresentation of underlying structure, the topology modification routines used during NanoWrap's shrink-wrapping iterations are quite conservative about when they will change the topology - i.e. they will only change topology if it is very obvious the current topology is wrong. They are also more reliable at removing spurious connections and holes than adding missing connections or holes. Because we bias the starting surface to be slightly outside the structure of interest, we are more likely to miss holes in areas of low density than to generate spurious holes. To assess the ability of our method to accurately capture membrane topology, we performed simulation of a synthetic structure with a range of hole sizes (Supplementary Figure S8). To avoid gaming the analysis we used an automated algorithm to set the threshold for the starting surface (see below). We also chose the curvature weight so as to minimise RMS error, not hole count. Under these conditions, the ability to capture a hole depends on a combination of localisation density, precision, and background level. At high densities and low levels of background, we can typically capture holes that are larger than around 3 times the localization precision, although we occasionally miss a hole we really should have captured. Manual tweaking of the parameters could have achieved improved the results. Correctly estimating surface genus is an area where we believe there is scope for improvement in our algorithm, especially in deciding when to modify the topology as we iterate.

#### Automatic threshold selection for starting surfaces

We expect that the starting surface should be close to the localizations used to generate it, and the median distance from each localization to the starting surface can be used as quality metric for the surface. By trying different thresholds within an optimisation routine (we use bisection) a threshold value leading to a good starting surface can be chosen. It practice it is helpful to aim for a distance (e.g. $-10$ to $-40$ nm) which puts the localizations just inside the surface and avoids the generation of spurious holes. This algorithm can be activated by selecting the 'Auto threshold' box in the Dual Marching Cubes settings, and will almost always generate good starting surfaces. Manual tweaking can, however, be useful if there are topological features (e.g. holes) which are right at the limit of being resolvable with the given localisation precision and densities.

#### Rational choice of curvature weighting

Similar to choosing a threshold for the starting surface, the distance between localizations and the reconstructed membrane can be a useful metric to determine whether an appropriate curvature weight has been selected. This time, however, we look at the distribution of distances rather than their median. Localizations are expected to scatter about the true membrane position due to their localisation error, with the distance between a localisation and the true position of that molecule expected to follow a scaled Chi(3) distribution. The choice of curvature weight is appropriate if the distribution of distances between localizations and the membrane is similar to that predicted by the localisation error. If the spread is larger, the curvature weight is too high, but if the spread is less than that predicted by localization error the algorithm is over-fitting to the points and the curvature weight should be increased. For performance reasons, we have not automated this, and setting the curvature weight remains a manual process. A function to plot the distance distributions is available under *Mesh->Show shrinkwrap residuals* within PYMEVis.

These distributions are also useful for showing if the number of iterations is suitable - if the residual distribution is heavily weighted to negative distances (inside the surface) this indicates that either more iterations are likely needed to allow the surface to converge.

### Supplementary Figures

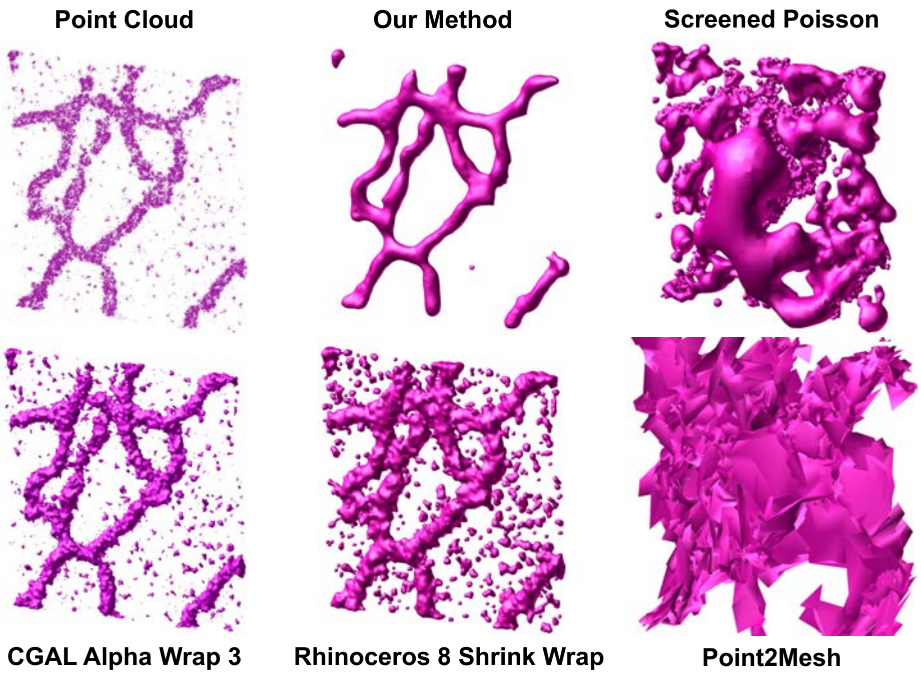


Figure S1: A comparison of our method to other common methods used to generate surfaces from a point cloud. Top left is an example point cloud from a subregion of an endoplasmic reticulum. A surface is generated from this point cloud with NanoWrap, Screened Poisson Reconstruction (18), CGAL's Alpha Wrap (22), Rhinoceros 8's shrink wrap algorithm (47) and Point2Mesh (23). NanoWrap allows the surface to adapt per-point based on localization positions and achieves a better representation of the underlying structure than other methods, which strictly adhere to the point cloud or generate a surface at a constant offset from each point. We manually experimented with the input hyperparameters of each algorithm to achieve the best surface in each case. A grid search may yield better surfaces for each of these methods, but we do not expect significant improvement, due to the nature of the input data. Both SPR and Point2Mesh require point normals as input. Point normals were estimated using Meshlab's (48) estimator. Note that these methods may perform well in the presence of true normals for a point cloud, but normal estimation usually relies on neighboring points, including noise points that confound the estimate. Point2Mesh requires the user to provide an initial starting surface. We provided it the same starting surface we used for our approach, which is a dual marching cubes (30) approximation on the density of the point cloud.

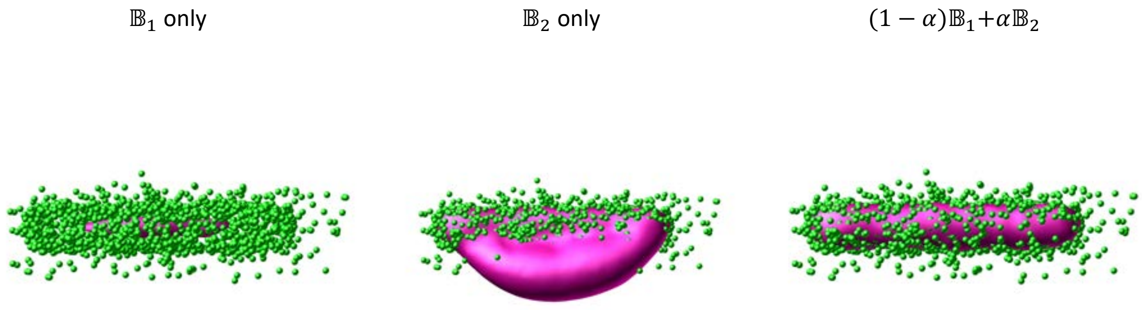


Figure S2: This figure demonstrates the independent influence of curvature forces $\mathbb{B}_1$ and $\mathbb{B}_2$ on the final NanoWrap surface. The localization point cloud is shown in green and the surface fitting these points is shown in magenta. $\mathbb{B}_1$ has area-minimizing tendencies and the surface pulls through the point cloud. $\mathbb{B}_2$'s preservation of area means that areas of the mesh far from point influence can establish stable "blebs" protruding from the true structure. The combination of forces subject to a point influence weighting $\alpha$ creates an accurate reconstruction of a surface. See the methods section for more details.

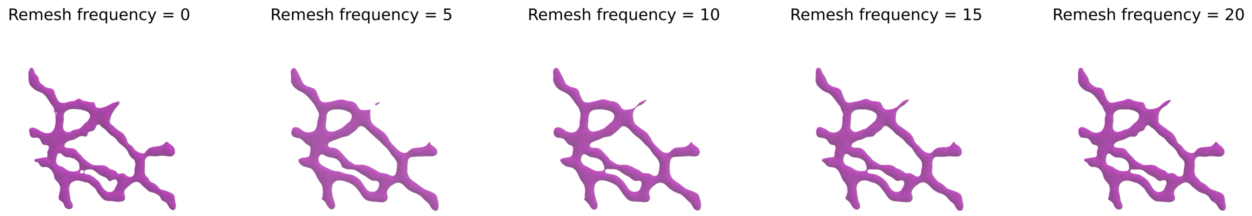


Figure S3: A NanoWrap surface fit over 39 iterations, remeshed every 0, 5, 10, 15, 20 iterations. The structure does not change significantly depending on remesh frequency.

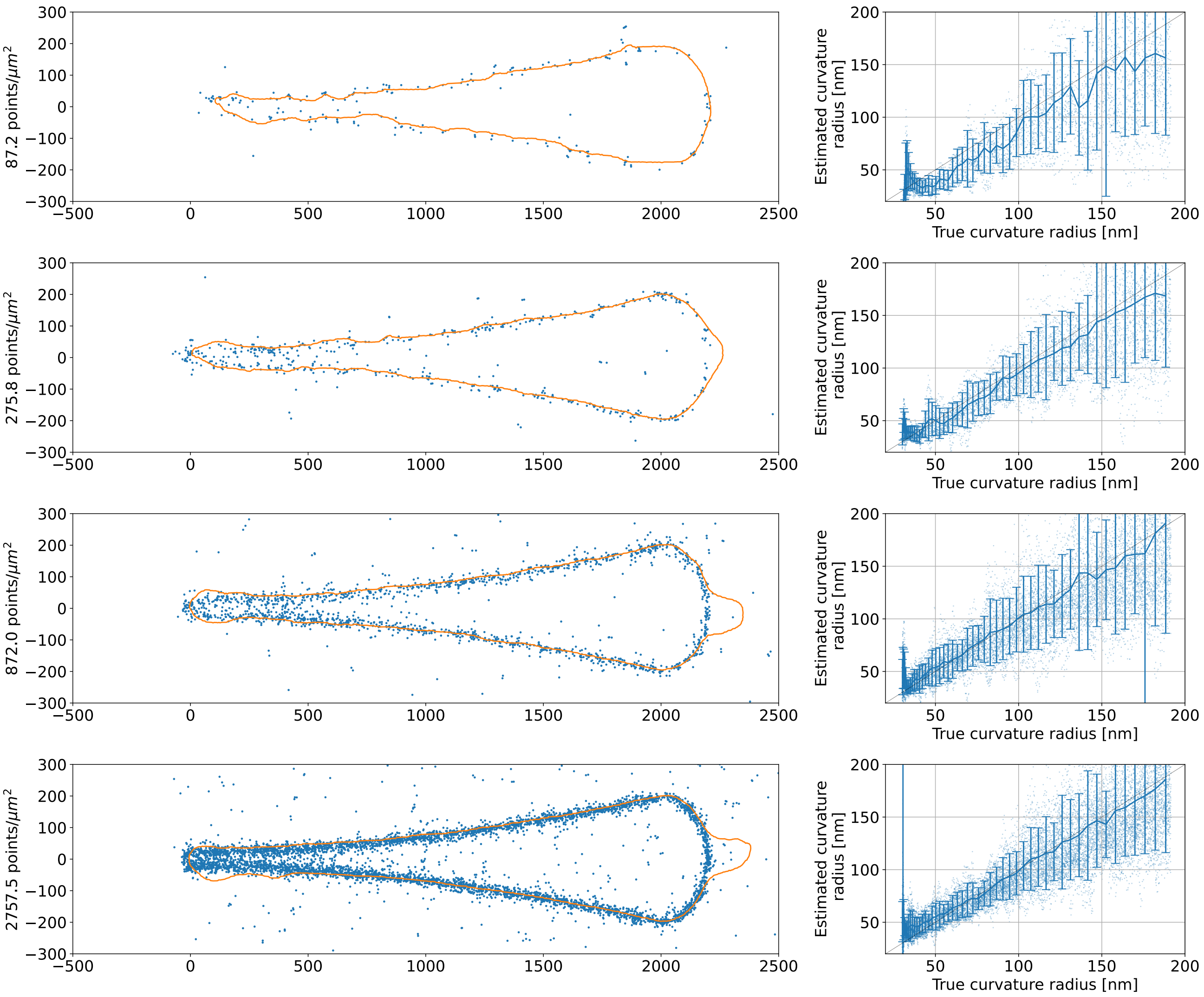


Figure S4: The radius of curvature of a reconstructed surface of a tapered cylinder as a function of density. The tapered cylinder's diameter scales between 30 and 200 nm over a 2000 nm distance as the square of the distance along the cylinder. The left column shows a cross section of the fit point cloud (blue) and the reconstructed surface (orange). The right column shows the estimated radius of curvature for each vertex of the fit surface along with an error bar indicating the full range of curvature estimates extracted from the vertices for each expected radius of curvature. As the density increases, the minimum radius of curvature that can be estimated decreases. The point cloud localization precision is fixed at 11 nm.

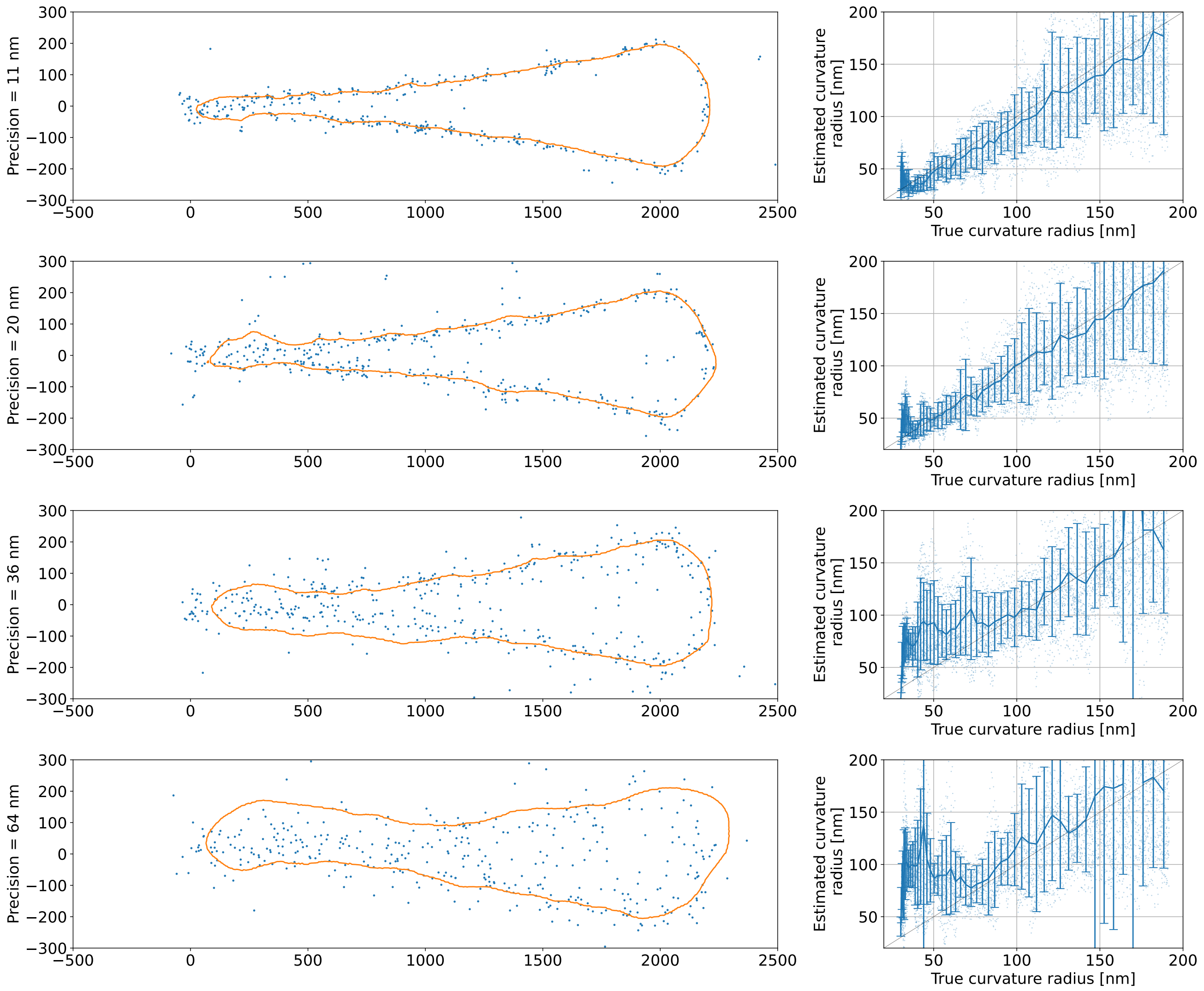


Figure S5: The radius of curvature of a reconstructed surface of a tapered cylinder as a function of localization precision. The tapered cylinder's diameter scales between 30 and 200 nm over a 2000 nm distance as the square of the distance along the cylinder. The left column shows a cross section of the fit point cloud (blue) and the reconstructed surface (orange). The right column shows the estimated radius of curvature for each vertex of the fit surface along with an error bar indicating the full range of curvature estimates extracted from the vertices for each expected radius of curvature. As the precision increases, the minimum radius of curvature that can be estimated scales to match. The point cloud density is fixed at 218 points $\mu m^{-2}$.

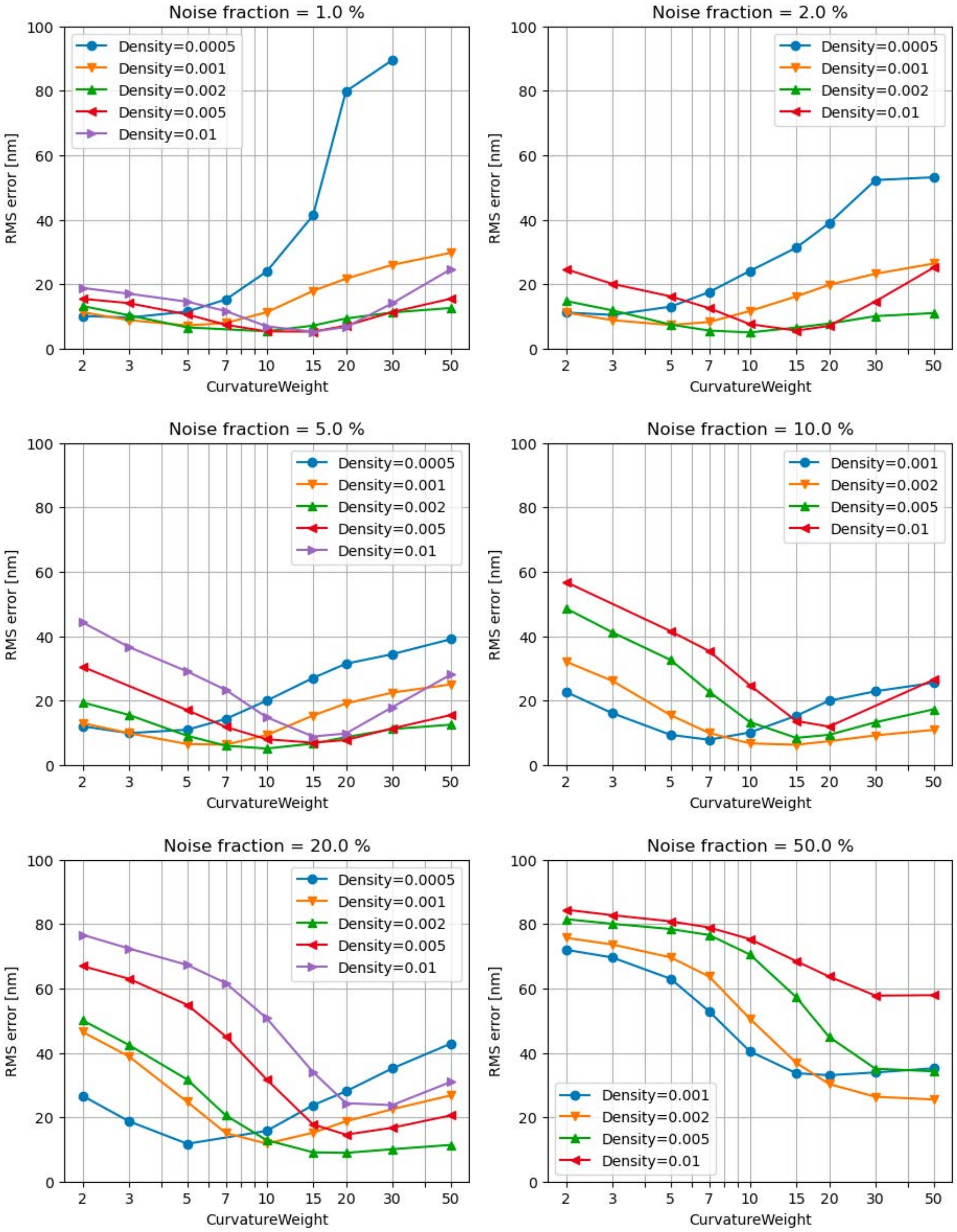


Figure S6: NanoWrap's sensitivity of fit performance to the chosen value of curvature weight at different values of structure density and background level. The optimal curvature weight increases with both increasing point density and background, but in each case there is a broad minimum.

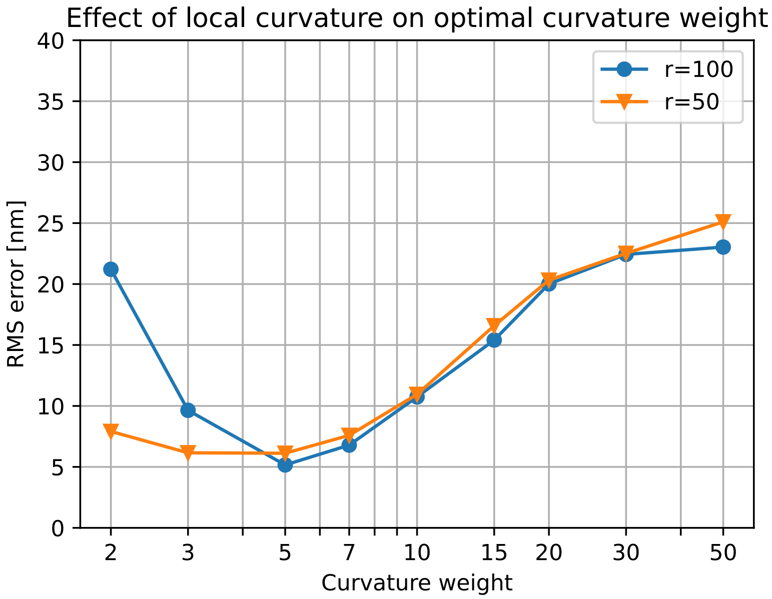


Figure S7: To test whether the optimal curvature weight depends on the underlying structure curvature we simulated a scaled version of the double-torus structure that had about twice the radius of curvature, whilst holding localisation precision, density, and noise levels constant in the middle of our range. There is only a very slight change in the optimal curvature weight, although the flatter structure shows a faster degradation in performance when the curvature weight is decreased below the optimum. This can be attributed to an additional regularising effect of the finite mesh size at small radii of curvature.

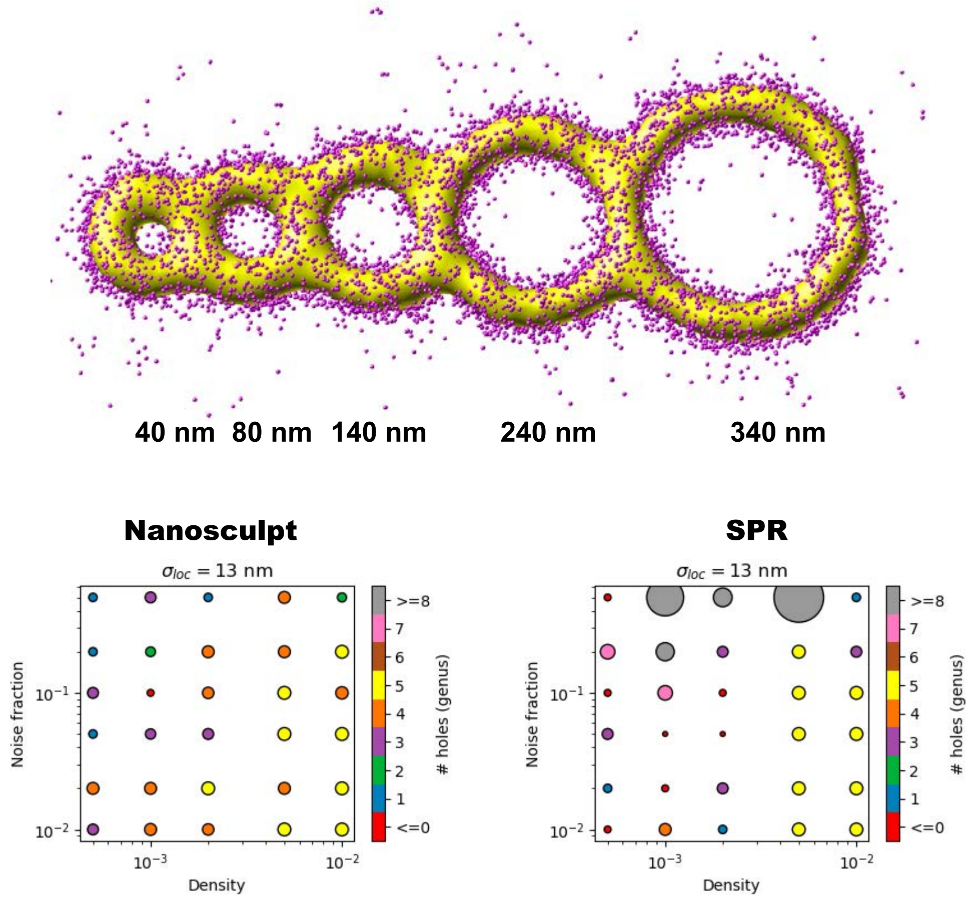


Figure S8: Ability of NanoWrap to correctly estimate the number of holes (genus) for a test structure using an automatically thresholded starting surface. Our ability to correctly estimate hole count is broadly comparable to SPR, but degrades a bit more gracefully at poor localization density. The genus estimation is very sensitive to the quality of the starting surface, and in the few cases where SPR correctly estimates the genus and we do not, the correct genus is easily obtained by manually tweaking the parameters for starting surface generation. This implies that there is still scope to improve the algorithm we use for intial surface estimation.

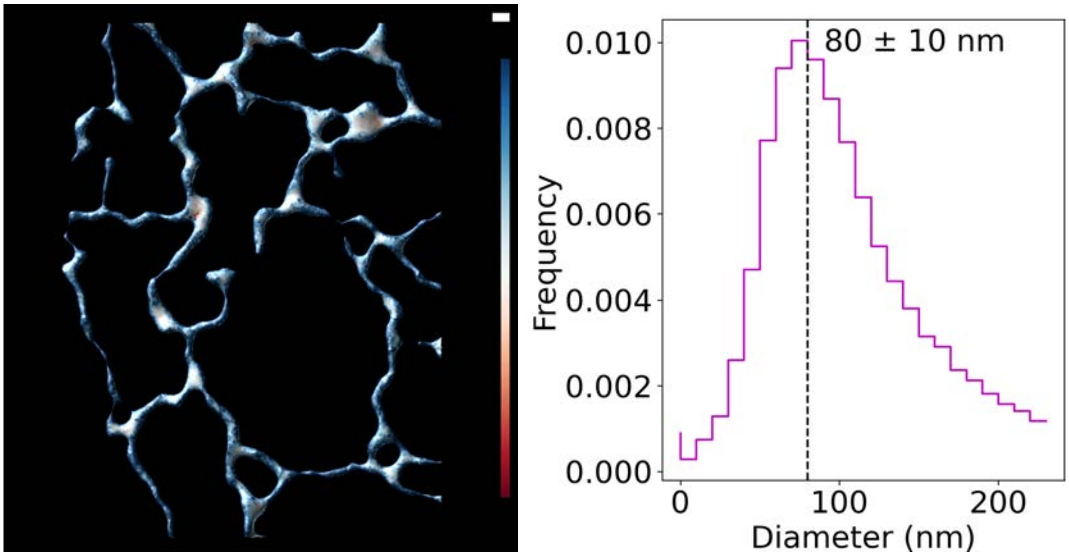


Figure S9: **Left,** Tubular ER from the lower left portion of Figure 3, colored by mean curvature. Lookup table: -0.01 to 0.01 $nm^{-1}$. Scale bar is 200 nm. **Right,** Histogram of $D = 2/k_{\mathrm{max},l}$ where $k_{\mathrm{max},l}$ is the maximal principal curvature of vertex $l$. The mean and standard error of the mean for the diameter distribution are reported.

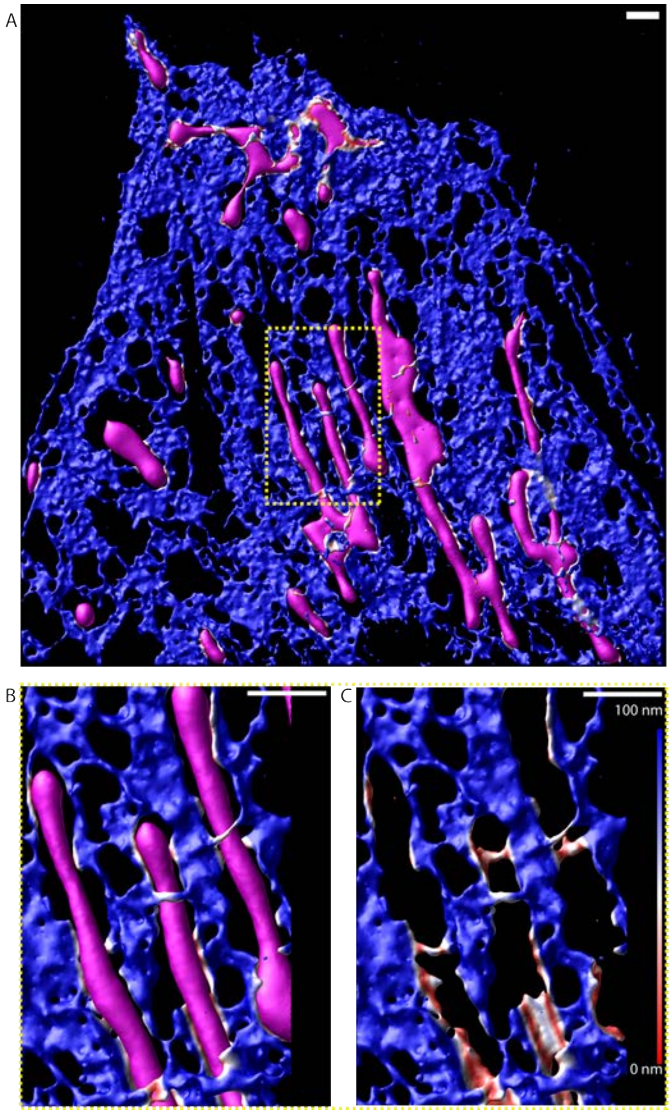


Figure S10: Distance between two surfaces. Surface generated from Sec61$\beta$ (ER membrane) localizations (red-white-blue) and surface generated from TOMM20 (outer mitochondrial membrane) localizations (magenta). The Sec61$\beta$ surface is colored by its distance from the TOMM20 surface (0 to 100 nm; red to blue). **A,** Full field of view, displaying both surfaces. **B,** Region of interest, showing both surfaces. **C,** The Sec61$\beta$ surface alone. Scale bars are 1 µm.

### USER GUIDE

This tutorial will walk you through using NanoWrap surface reconstruction within PYMEVisualize to create 3D surfaces based on a provided 3D point-cloud data set (`user-guide-data.hdf`). After completing this tutorial, you should be able to apply the same approach to fit your own 3D point-cloud data sets.

If you are unfamiliar with using PYMEVisualize, we strongly encourage you to first install PYME (see `https://python-microscopy.org/doc/Installation/Installation.html`) and go through the PYMEVisualize User Guide (`https://python-microscopy.org/doc/PYMEVis.html`) to understand the basics of the program before moving on to this tutorial (27).

If using the executable installers (recommended), the NanoWrap plugin should be automatically installed alongside PYMEVis. If this does not work, or for platforms where an executable installer is not available (currently linux and mac M1 native), PYME's installation instructions also cover how to set up Anaconda (`https://anaconda.org/`) and the Anaconda PYME environment. The NanoWrap plugin will then need to be installed manually following the instructions at `https://github.com/python-microscopy/ch-shrinkwrap`.

#### Verify Plugin Installation

Launch PYMEVis by either selecting from the start menu (windows) or by typing `PYMEVis` at the console (mac, linux) and hitting enter and confirm there is the menu option **Mesh → Shrinkwrap membrane surface** . Depending on how PYME was set up (executable vs manual conda-install) you might need to activate the conda environment - `conda activate PYME` - before the `PYMEVis` command is available. If the menu option is not visible, recheck the installation instructions at `https://github.com/python-microscopy/ch-shrinkwrap`.

#### Download and import tutorial data set

Download the data set `user-guide-data.hdf` found in the supplementary materials. Launch a PYMEVisualize window by entering `PYMEVis` into an Anaconda prompt with the PYME environment activated. Navigate to **File → Open** and select the `user-guide-data.hdf` file. To improve the visualization of the data, under the "Layers" side tab change "Method" to *pointsprites*, "Colour" to *z*, "Alpha" to *0.2*, and "Point size" to *10*. Your PYMEVisualize window should look identical to Figure S11 now.

#### Create initial isosurface

You can learn about PYMEVisualize isosurfaces and how to make them here. Once you're familiar with how to create an isosurface, use the parameters shown in Figure S12 to create an isosurface based on the `user-guide-data.hdf` points. Note that you can alternatively check "Auto threshold" to let NanoWrap estimate the ideal threshold density of the starting surface. See Supplementary Materials for a description of how automatic thresholding works. Once finished, you should see a 3D surface rendered with your points. Change "Method" to *wireframe* to more easily visualize how well the isosurface adheres to the point cloud. It should look something like Figure S13. At this point, it is a good idea to assess the overall accuracy of this initial isosurface. You can alter the parameters for the isosurface in the **Data Pipeline** tab on the left under **DualMarchingCubes**. It is typical to test several parameters before settling on the best options with "N points min" and "Threshold density" being the two major parameters requiring tweaking. It should be noted that this data set is a small cropped region of interest (ROI) from a much larger data set. It is highly recommended that you test parameters for the initial isosurface (and the fitted surface) on a small ROI of your data that has enough points to test the structure you're trying to create a surface from, but no more than that (about 20,000 points is ideal). The time required to create the surfaces scales roughly with the number of localizations, so testing parameters on large data sets is impractical. Once you establish which parameters are best with your smaller ROI, you can apply the same ones to the full data set. Learn how to create an ROI in PYMEVisualize here.

#### Fit surface with NanoWrap

Now that we have our initial isosurface, we can use it to create a fitted surface using NanoWrap. You will find this option under **Mesh → Shrinkwrap membrane surface**. The first window that pops up is requesting parameters to create an isosurface. If you haven't done so already make sure to create an isosurface because it is required for fitting. If you already have an isosurface, which you should if you are following this user guide, you can close this first window. The next window that pops up is requesting parameters you'd like to use for fitting. Refer to Table S2 for descriptions of what each parameter is and what typical values are for it. For this tutorial, use the parameters shown in S14 or check "Auto threshold". This process can take a minute or so. Check the Anaconda prompt for output to confirm it is running. Once finished, your surface should look similar to S15.

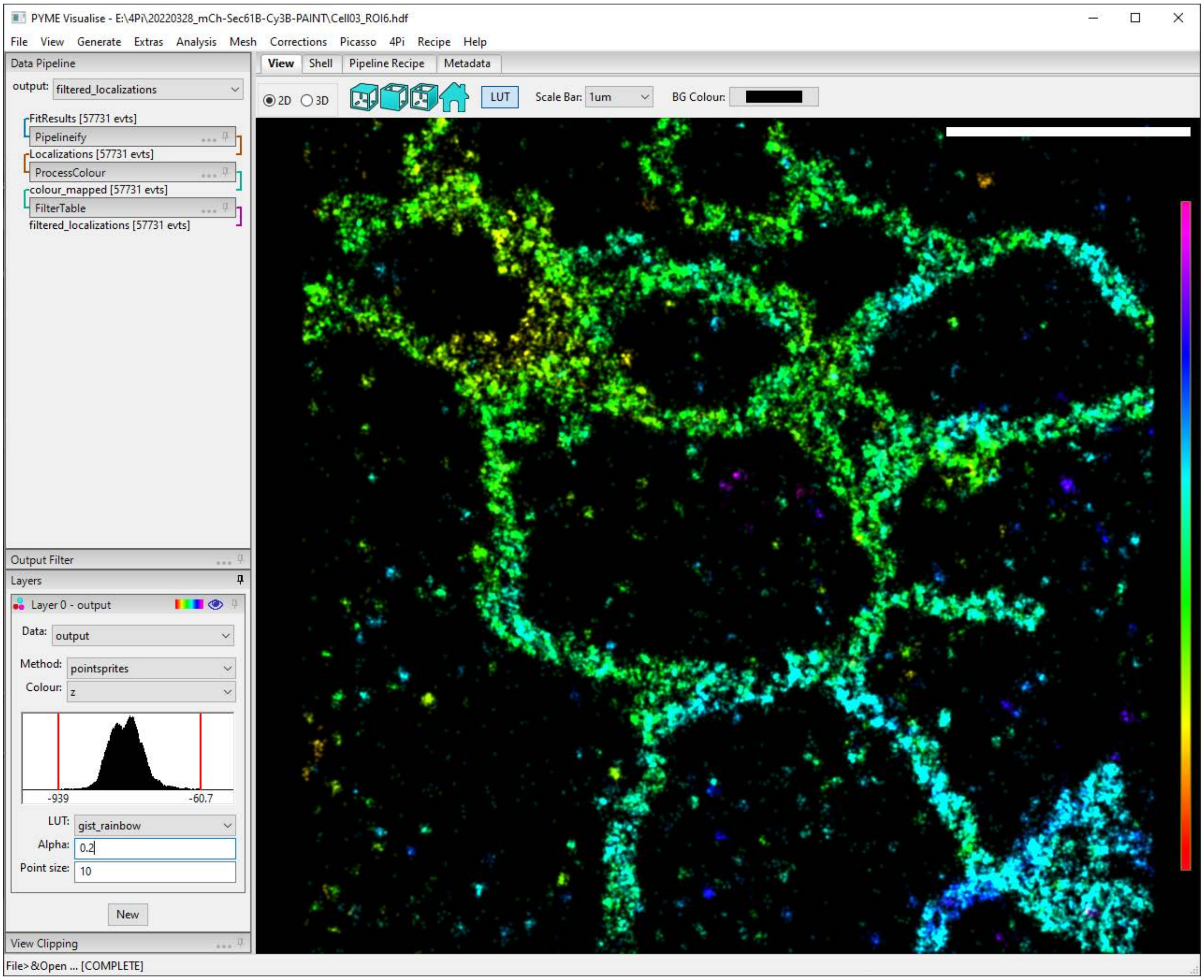


Figure S11: Data from `user-guide-data.hdf` displayed as 10 nm point sprites with 0.2 alpha. The points are colored by their location in the z-direction according to the lookup table on the right.

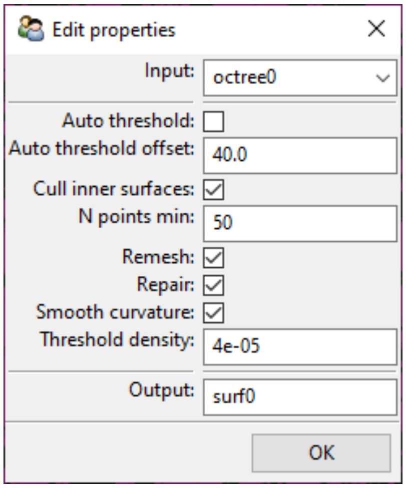


Figure S12: Ideal parameters to use to create an isosurface based on `user-guide-data.hdf`.

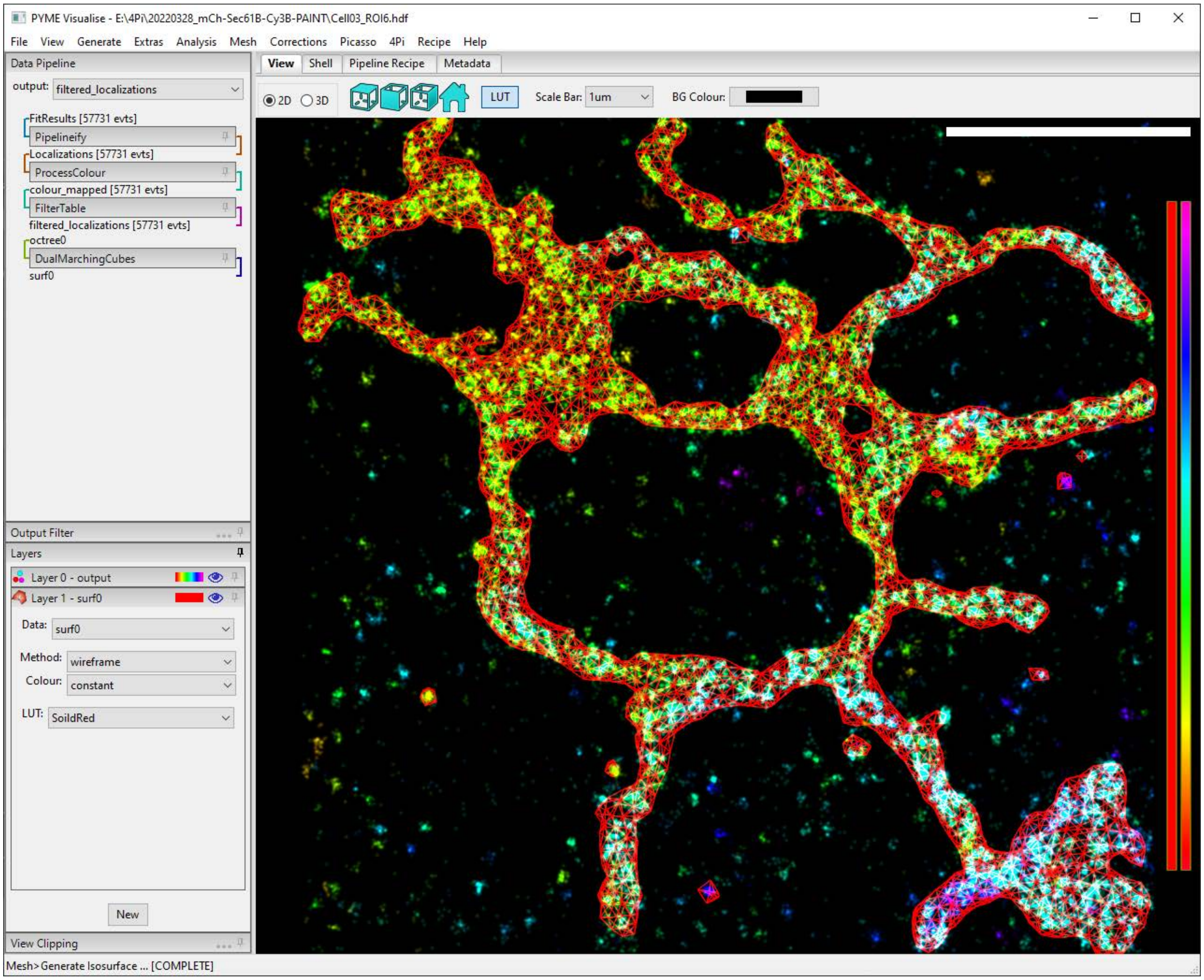


Figure S13: The isosurface generated using the parameters in S12. The isosurface is displayed using the *wireframe* method to make it easy to assess how well it fits the point cloud.

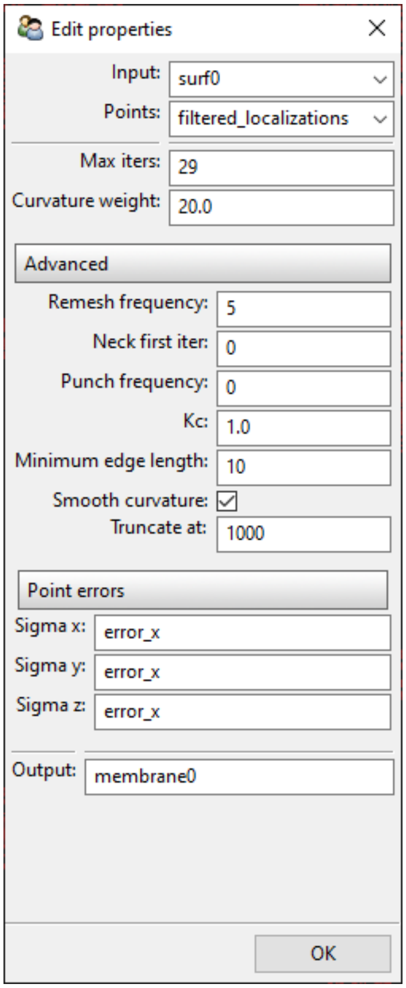


Figure S14: Ideal parameters to create a surface based on `user-guide-data.hdf`.

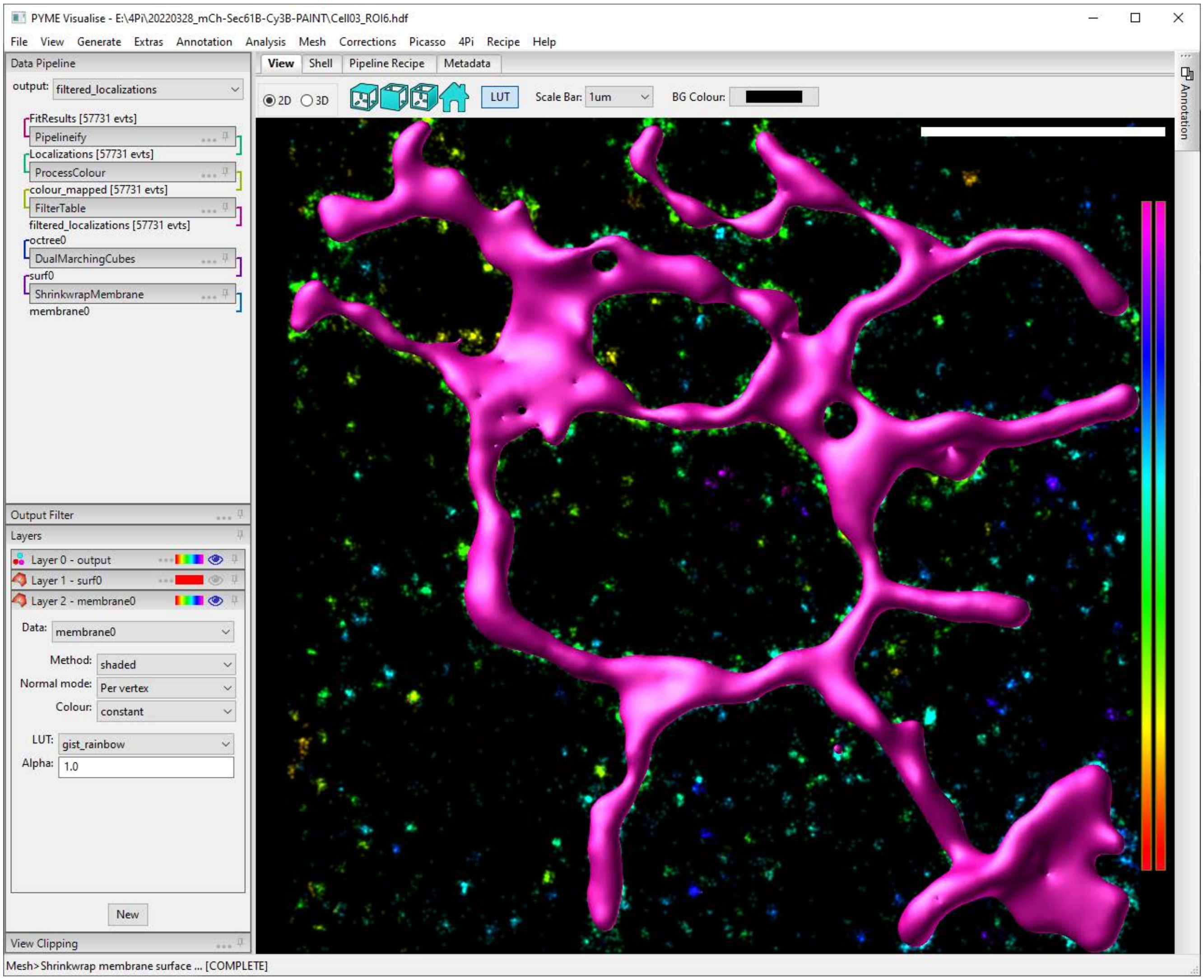


Figure S15: The surface generated using the parameters in S14.

| Parameter | Description | Standard values |
|---|---|---|
| input | The data source containing the coarse isosurface. | `surf0` |
| Points | The data source containing the points to fit. | `filter_localizations` |
| Max iters | Maximum number of fitting iterations. | 10 - 100 |
| Curvature weight | The contribution of curvature (vs. point attraction force) to the fitting procedure. Higher values create smoother surfaces. | 10 - 100 |
| Remesh frequency | Remesh the isourface every N iterations. Helps keep the fitting numerically stable. Should be often. | 5 |
| Neck first iter | Every neck first iter iterations, check for and remove necks in the mesh. | 9 (0 to not use necking) |
| neck_threshold_low | Vertices with Gaussian curvature below this threshold are necks. | -1.00E+03 |
| neck_threshold_high | Vertices with Gaussian curvature above this threshold are necks. | 1.00E+02 |
| Punch frequency | Every punch frequency iterations, check for and add holes in regions of the mesh where there is a continuous empty area in between two "sides" of the mesh. | 0 |
| Kc | Lipid stiffness coefficient of membrane in eV (can be looked up in the literature). | 0 - 1 (20k_bT) |
| Minimum edge length | Small length that the edges joining surface vertices can be. Smaller means the surface is more finely sampled. Setting it to 10 is usually sufficient. Setting it to -1 removes any limit. | 10 |
| Smooth curvature | Replace the fit curvature of a vertex with the average curvature of it and its neighbors | `True` |
| Truncate at | Stop after this many iterations no matter what. Useful for visualizing the behavior of a surface over time | 1000 |
| error_x | The variable in the points data source containing localization precision in the x-direction. If error is only known for one axis, supply it here and it will be assumed for all axes. | |
| error_y | The variable in the points data source containing localization precision in the y-direction. | |
| error_z | The variable in the points data source containing localization precision in the z-direction. | |
| output | The name of the data source that will contain the fit isosurface. | `membrane0` |

Table S2: Descriptions and standard values for all parameters used in this paper's algorithm.

### Additional tips for fitting surfaces

The following are tips for situations that did not come up in the tutorial, but are worth mentioning:

- An error that can occur when attempting to fit a surface is the "singular matrix" error. This simply means that the operation reached a point that was mathematically impossible to calculate. It is most often caused by several surfaces being present after creating an isosurface, especially small surfaces that are often capturing background localizations. Most of the time, this can be fixed be removing all but the largest surface present. To do so, enter the following line the the **Shell** tab in PYMEVisualize: `pipeline.dataSources['surf0'].keep_largest_connected_component()`.

NOTE `surf0` should be the name of **your** isosurface. By default it is `surf0`, but can be a different name for various reasons.

NOTE This solution clearly isn't always reasonable to use. For data from an ER protein, it is reasonable since the ER is continuous. However, data from a mitochondrial protein is not so reasonable since mitochondria are discontinuous. In the latter case, try to change the initial isosurface parameters, especially *N points min* and *Threshold density*, to resolve the error when fitting.

- If attempting to create several separate fit surfaces from multi-color data, make sure you are fitting to only the points belonging to the desired channel. By default, the algorithm uses `filtered_localizations` which includes all points in the data. To learn how to extract color channels from multi-color data in PYMEVisualize, please read the relevant documentation here.
